## Supplementary Material for "Single-cell motion of magnetotactic bacteria in microfluidic confinement: interplay between surface interaction and magnetic torque"

‡ first authors

* corresponding author

1 Biomaterials department, Max Planck Institute of Colloids and Interfaces, Potsdam, Germany
2 Theory and Bio-systems department, Max Planck Institute of Colloids and Interfaces, Potsdam, Germany
3 Biological physics and Morphogenesis group, Max Planck Institute of Dynamics and Self-Organization, Göttingen, Germany
4 Aix Marseille Université, CNRS, CEA, BIAM, 13108 Saint Paul lez Durance, France
5 Institute for the Dynamics of Complex Systems, University of Göttingen, Göttingen, Germany
6 Department of Physics, Institute for Advanced Studies in Basic Sciences (IASBS), Zanjan 45137-66731, Iran

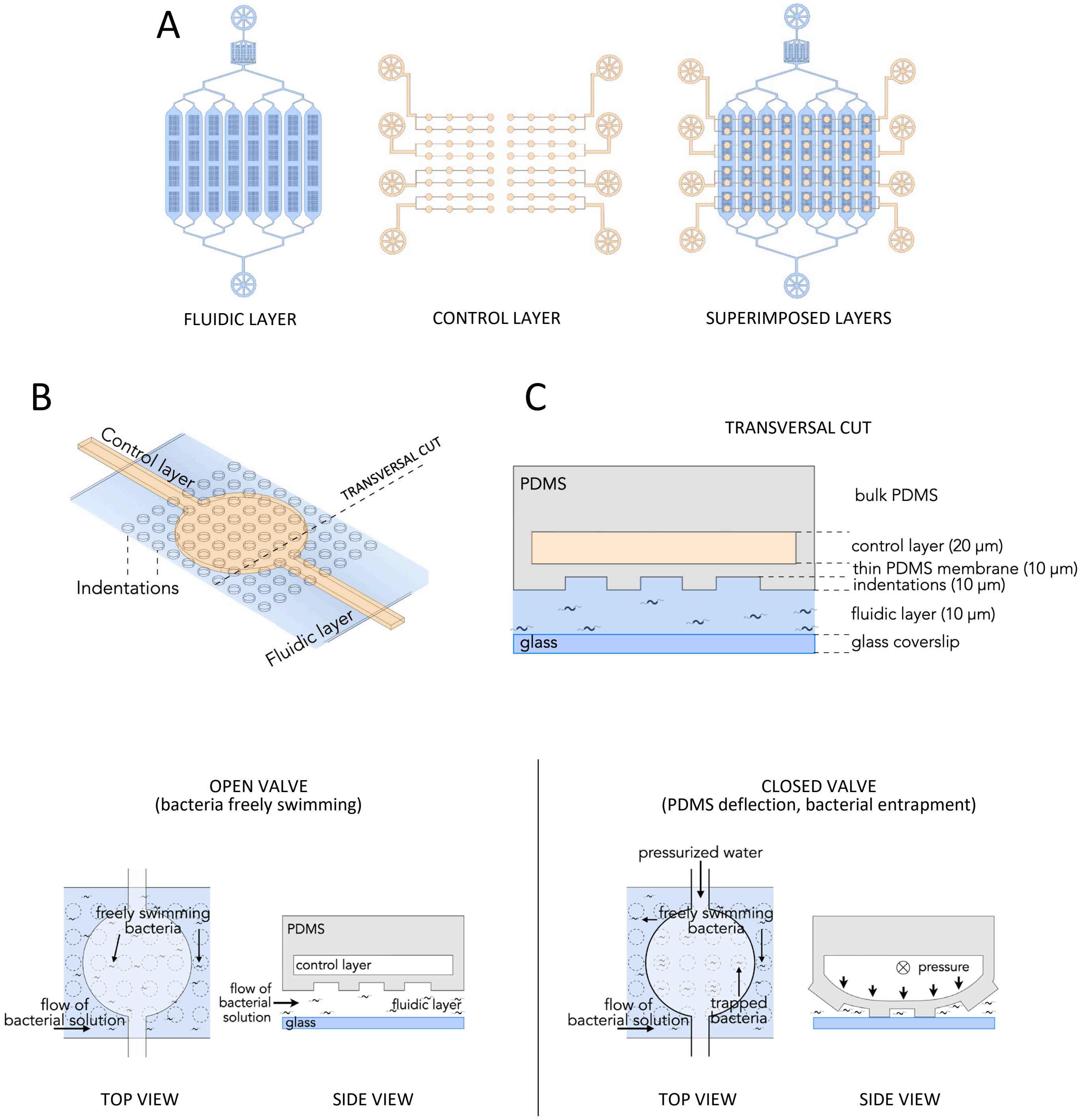


**Supplementary Figure 1. Schematics of the device configuration** A) Sketches of the fluidic (blue) and control (cream) layers and their superimposition after device assembly. The fluidic layer features one inlet and one outlet, whereas the control layer features eight inlets and no outlets. B) Zoomed 3D sketch showing how the control layer (cream) and the fluidic layer (blue) are perpendicularly positioned with respect to each other. The control layer contains circular valves which lie directly on top of an array of indentations present on the ceiling of the fluidic layer. Each microsystem features 64 valves with the possibility to generate up to 12 sealed microtraps per valve. (C) Transversal view of a cut in the microsystem (bold dashed line in B) showing the architecture of the control layer, PDMS membrane, indentations, and fluidic layer. The colour code is the same as in A and B. Note that the sketch has not been made to scale. (D) Top and side views showing the actuation mechanism leading to the microtraps' closing. When pressure is applied on the control layer, the water that fills the microchannels is pushed and deflects the PDMS membrane. When the pressure is high enough, the PDMS membrane deformation causes the indentations to come into contact with the glass bottom, generating a closed volume and trapping any bacteria within.

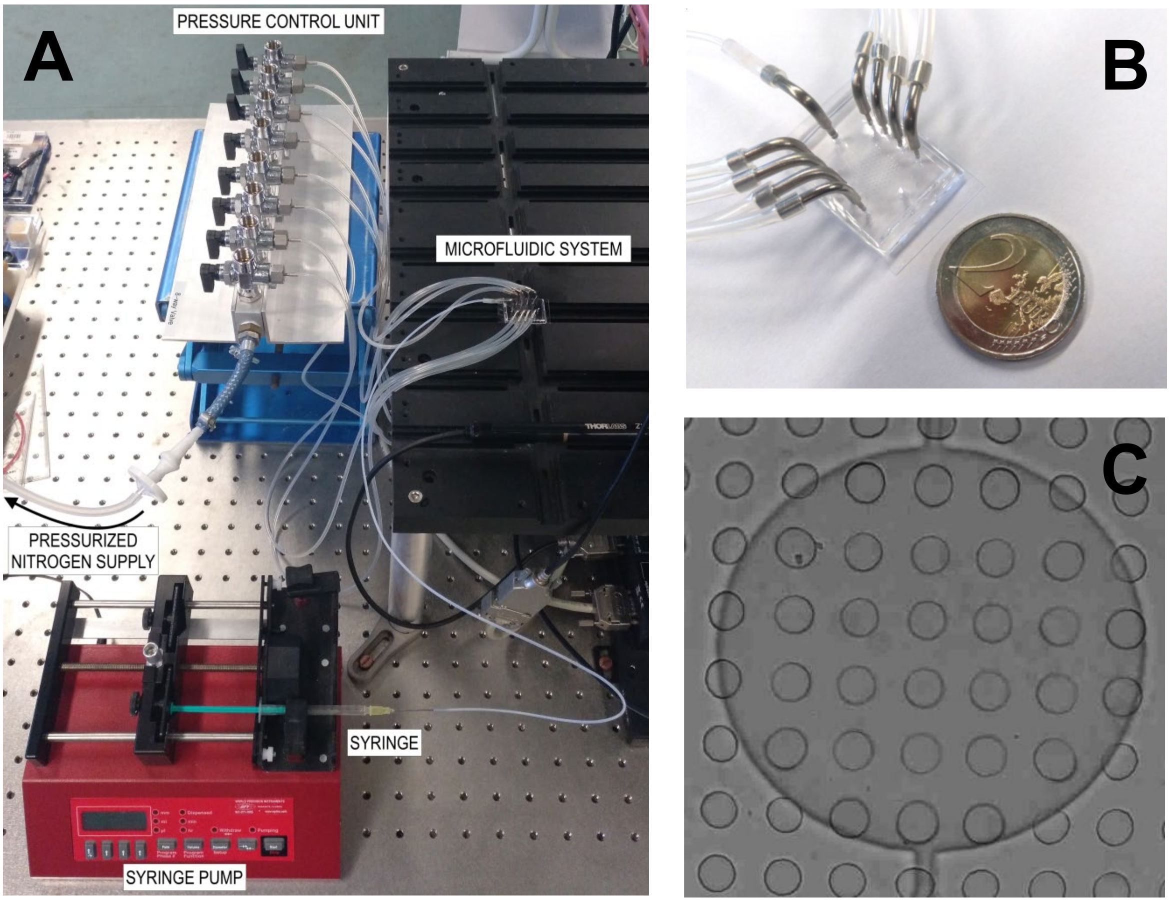


**Supplementary Figure 2. Microfluidic chip.** A) Microfluidic set-up: the PDMS microfluidic system is connected by of the fluidic layer to the syringe, and from the control layer to the pressurized nitrogen supply. B) Connections of the microfluidic system with its connections to the fluidic layer (metal connector in the center) and to the control layer (metal connectors on the sides). C) Bright field image of a control valve (big circle) on top of an array of microtraps (small circles). As a reference, each microwell is 35 µm in diameter.

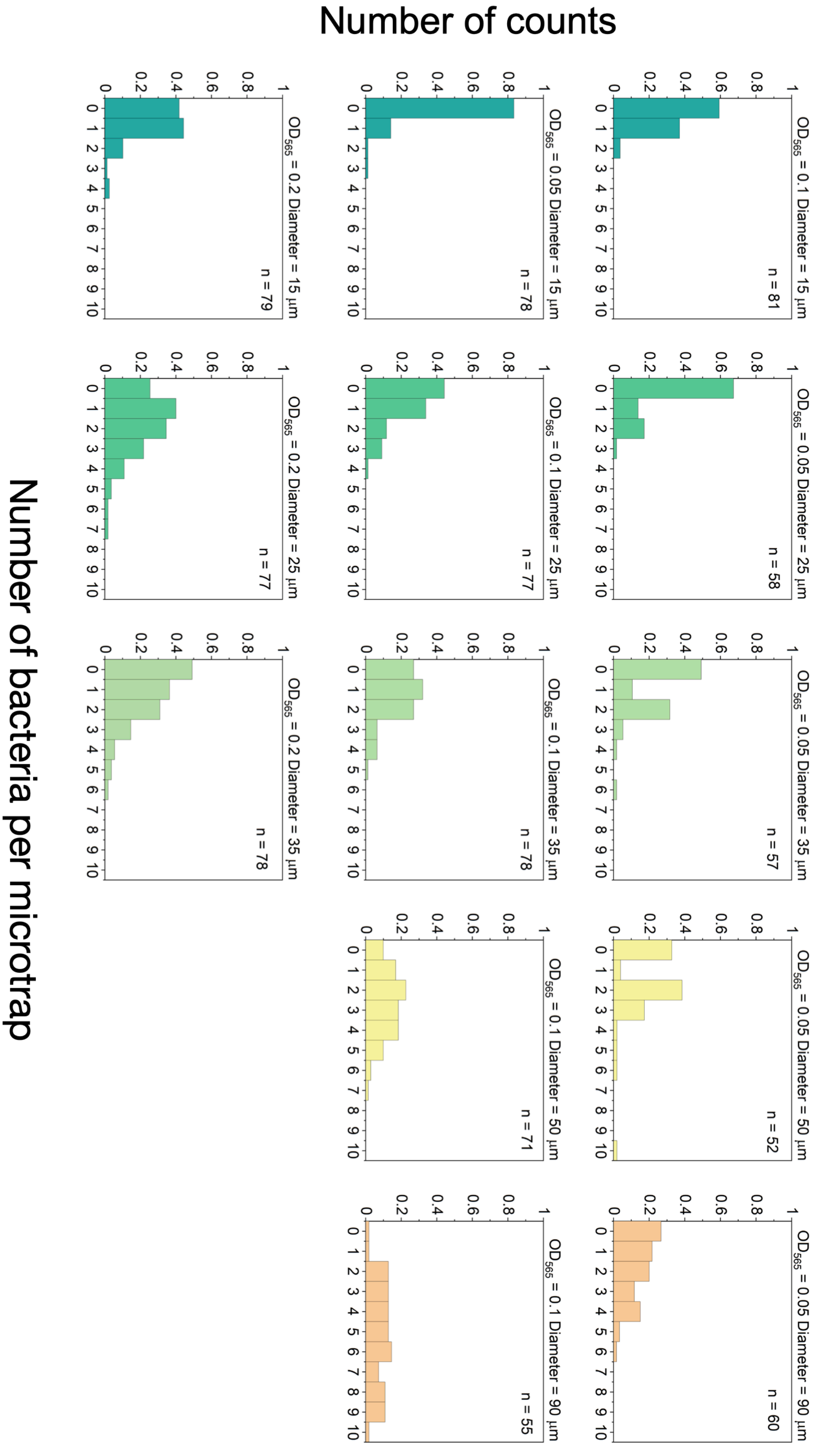


**Supplementary Figure 3. Number of bacteria in the microtraps.** Histogram of the number of bacteria per trap for three different optical densities (OD=0.05, 0.1 and 0.2) for each microtrap diameter at wavelength 565 nm. For $OD_{565}$ = 0.2, the bacteria within the biggest microtraps were not counted as it was difficult to determine the exact number due to their high number inside the microtrap.

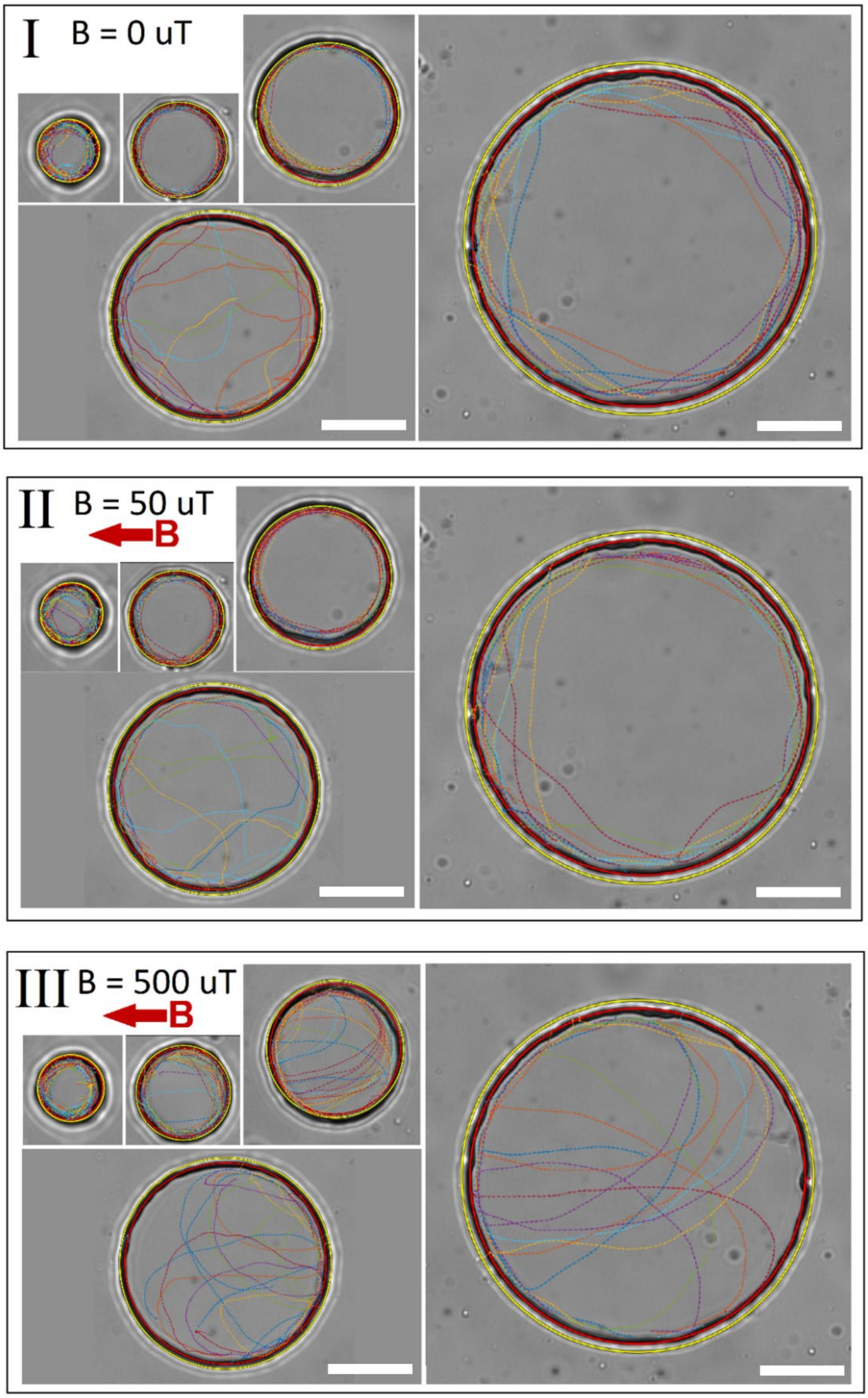


**Supplementary Figure 4. Bacteria trajectories in traps of different sizes**. Panels I, II, and III show the trajectories in 0, 50, 500 $\mu$T, respectively. Different trap sizes with radii of 7.5, 12.5, 17.5, 25, and 45 $\mu$m were examined. The red circles are the fitted circles to the trap wall and determine the boundary of the trap. The yellow circles are cutoffs with radius 1.1 times the trap radius, used to avoid considering any unwanted erroneous tracking points outside the trap. Scale bars are 20 $\mu$m.

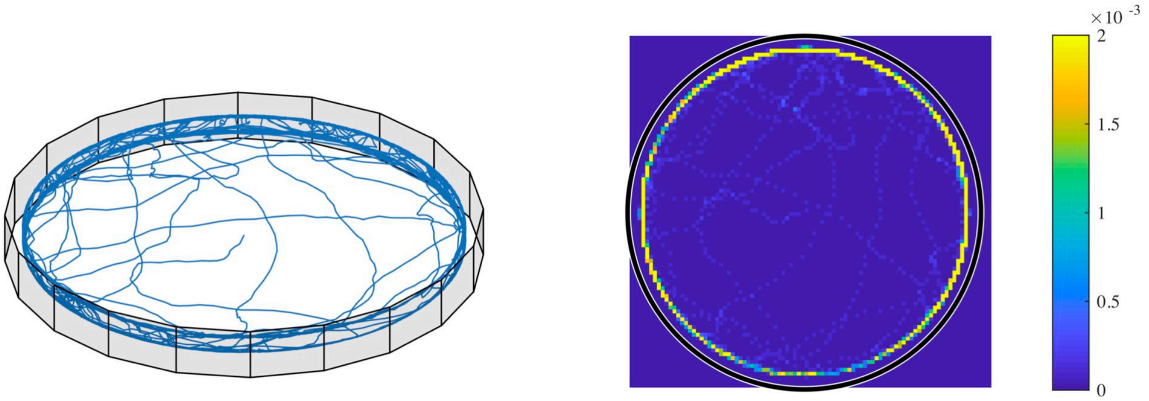


**Supplementary Figure 5. Simulated 3D trajectories without wall torque.** 3D trajectories and heat-map for the simulated case of trap radius 45 $\mu$m, with $A_w$= 6 but $T_w$ = 0. It can be clearly seen that the bacteria accumulate excessively at the border due to the wall torque set to 0 (heat-map scale saturated at 0.002; the color bar shows the normalized counts).

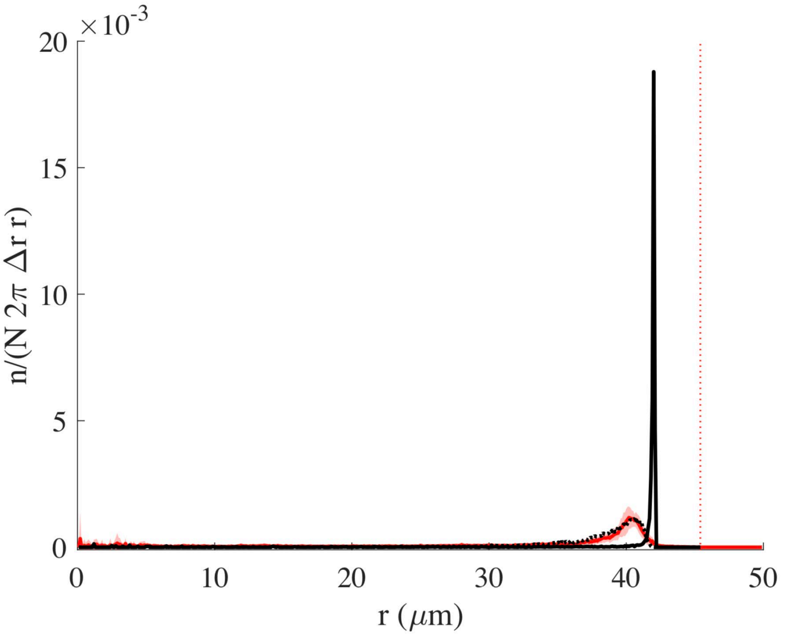


**Supplementary Figure 6. Radial distribution without wall torque.** The distribution corresponding to Supplementary Figure 5 (with $A_w$= 6 but $T_w$ = 0) can be seen here in black, together with the experimental distribution in red and with the matching simulation curve obtained with $T_w$=2.8 (black dotted curve). The vertical red line represents the wall.

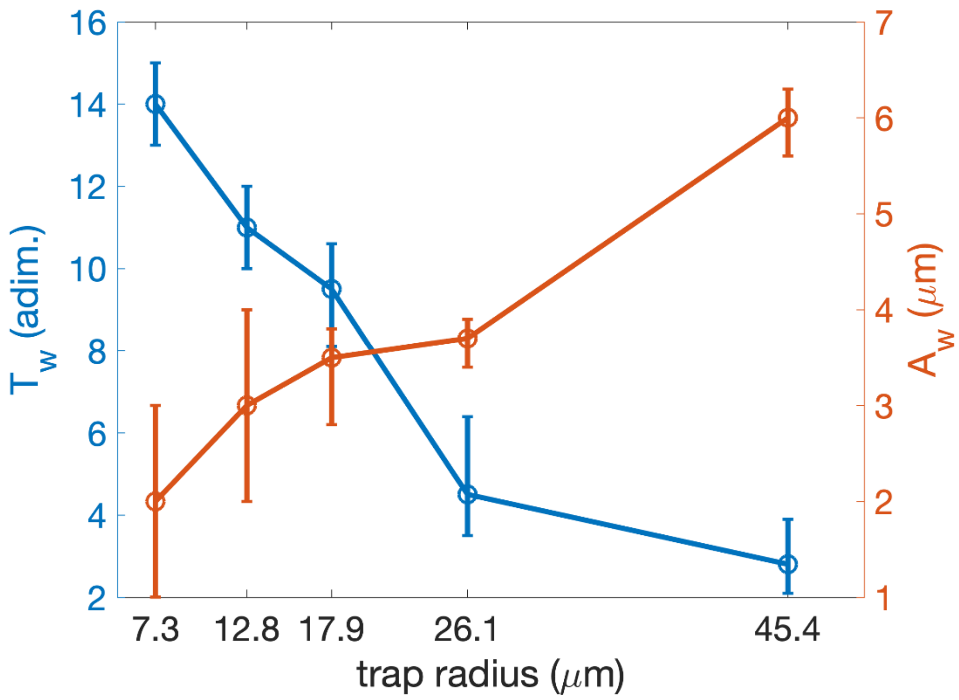


**Supplementary Figure 7. $T_w$ and $A_w$ as function of the trap radius.** Dependence of the parameters $T_w$ and $A_w$ (which provided closest data to the experiment minimising the adjusted R squared, see Table 3 of the supplementary information) on the measured trap radius, with the relative errors calculated as explained in Supplementary Table 3.

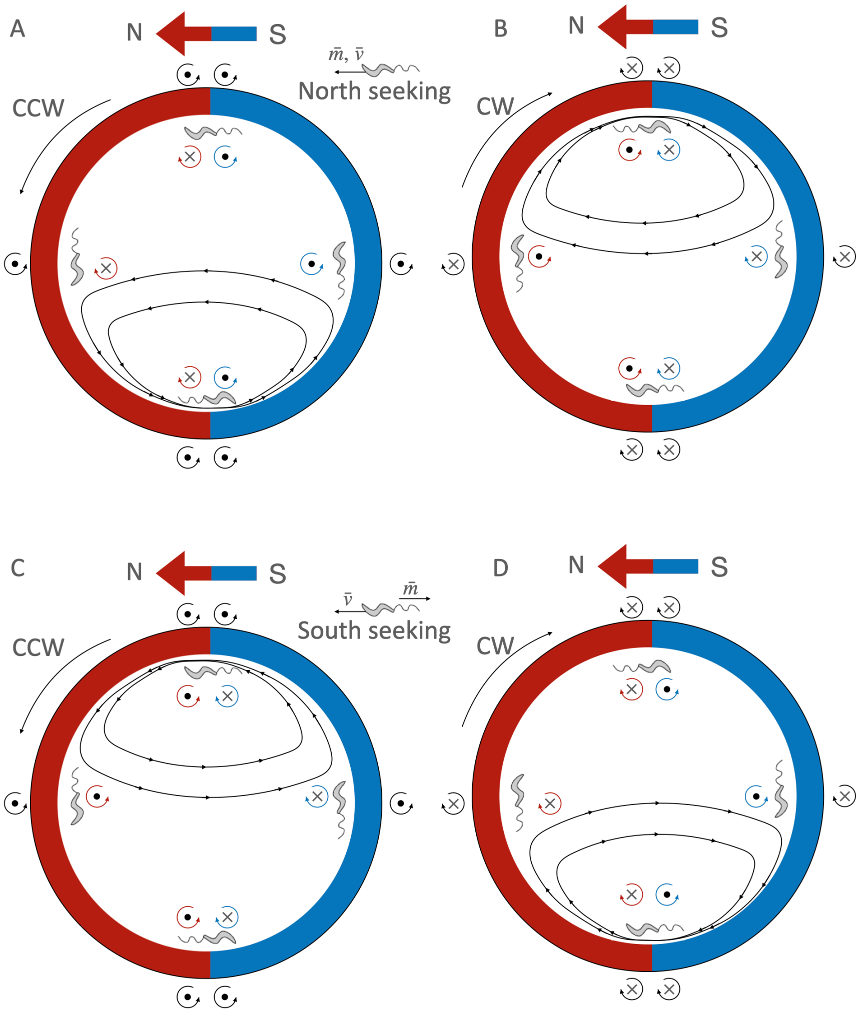


**Supplementary Figure 8. Interplay between wall-interaction torque and magnetic torque.** A) For a north-seeking bacterium (with the magnetic moment aligned with the velocity of the bacterium, thus swimming with the north of its magnet in front) swimming counterclockwise (CCW); B) north.-seeker swimming clockwise (CW); C) for a south-seeker (with the magnetic moment opposite to the velocity vector, swimming with the south of its magnet in front) CCW; and D) for a south-seeker CW. The red color indicates the north of the external magnetic field, and the blue the south. The blue/red arrow show the external magnetic field direction. The circular arrows in black indicate the direction of the wall torque; in red, the magnetic torque in the north part of the trap, and in blue in the south part. The dot at the center of the torque arrow indicates that the torque vector points out of the image, and the cross that it points into the image. Inside the trap, the black lines with arrows indicate the trajectories of the bacteria. Lower velocities and higher magnetic moments correspond to trajectories with smaller radii, and vice versa, higher velocities and smaller magnetic moments correspond to the trajectories with bigger radii. While the sign of the wall torque is given just by the swimming direction (CCW or CW), the sign of the magnetic moment depends on the position of the bacterium (if it is in the north side or south side of the trap), and on the orientation of the magnetic moment with respect to its velocity. When the wall torque and the magnetic torque point in the same direction, they sum up and push the bacterium off the wall and into the middle of the trap, producing the U-turns.

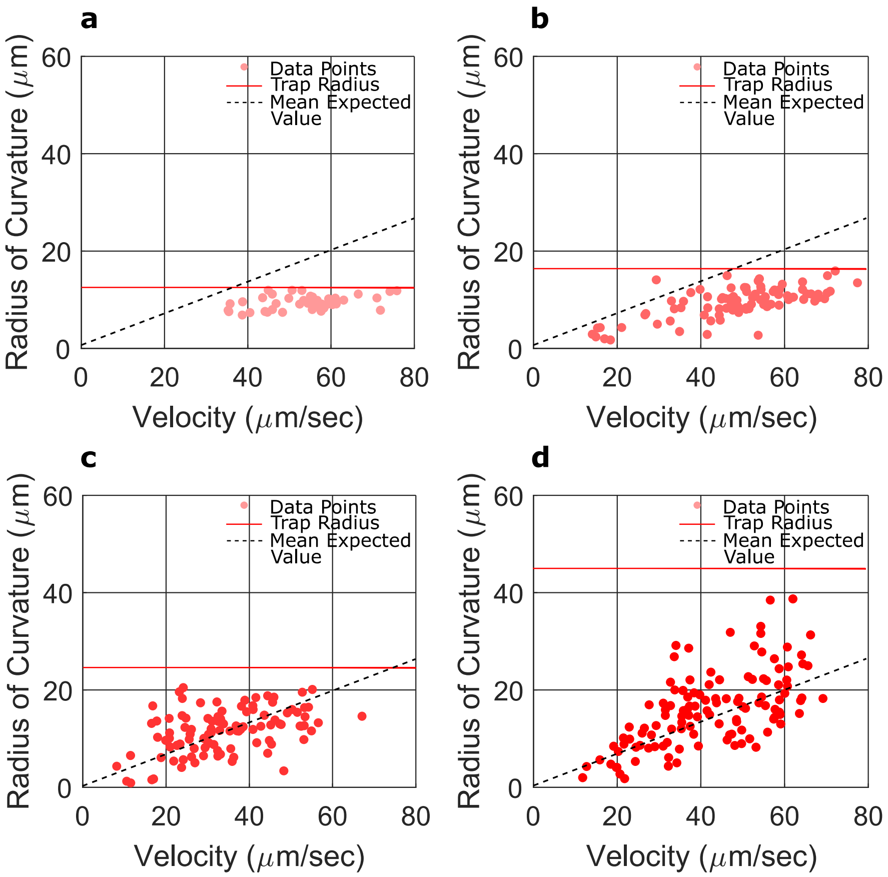


**Supplementary Figure 9: Radii of U-turns vs velocities for different microtrap sizes.** (a-d) The red dots show the U-turns radii for the microtrap sizes with radii of 12.5, 17.5, 25 and 45 µm, respectively. No U-turns were observed in the smallest microtrap size with radius of 7.5 µm. The horizontal red solid line represents the radius of the corresponding microtrap, and the dashed lines show the expected radii according to the mean value of the magnetic moment extracted from TEM images (see the Methods section for more information). As shown in the figures, for smaller microtrap sizes (radius of 12.5 µm and some part of trap with radius of 17.5 µm), the small microtrap radius avoids U-turns to happen normally and limits their appearance. Therefore, a threshold radius can be defined for a microtrap (~ $\boldsymbol{R_{Tresh}}$ = 22.5 µm, according to these data), for which in smaller sizes the bacteria would not perform U-turns properly and the surface interactions overcome the tendency of bacteria to do them. However, for the bigger microtrap sizes, for which the radius of the microtraps is greater than the overall radii of the U-turns, the bacteria can do U-turns freely.

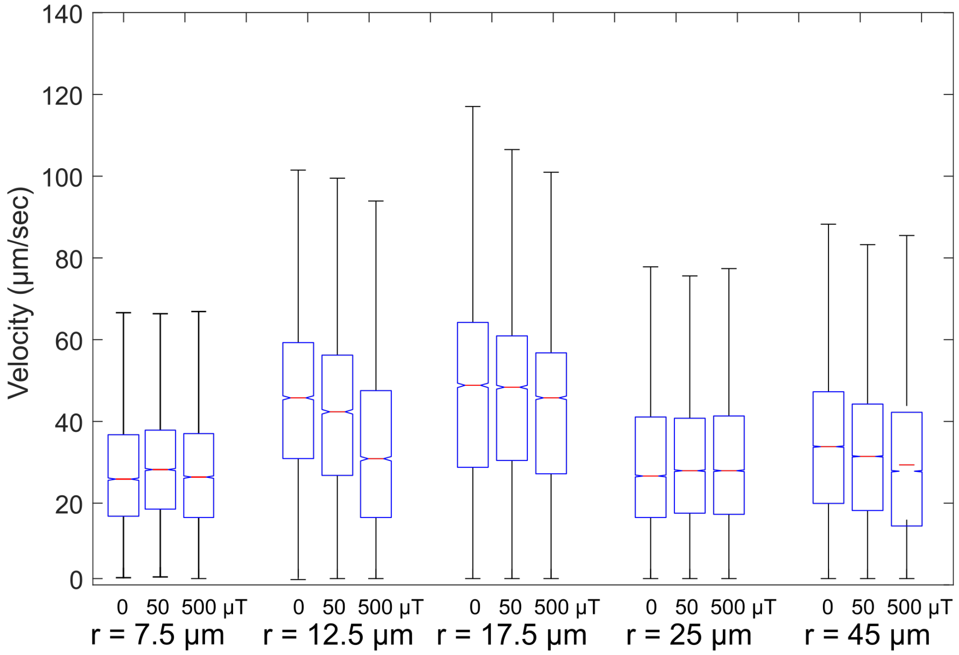


**Supplementary Figure 10: Velocity distributions.** Velocity distribution in different trap sizes and different magnetic fields. As shown in the above box plots, different bacteria have a wide range of velocities.

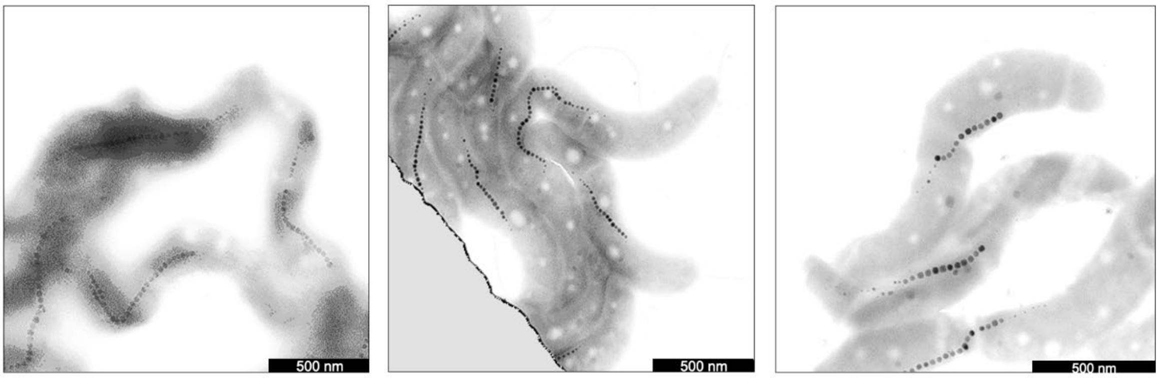


**Supplementary Figure 11. Magnetosomes TEM images.** Selected TEM images showing MSR-1 bacteria with magnetosome chains. Scale bars are 500 nm.

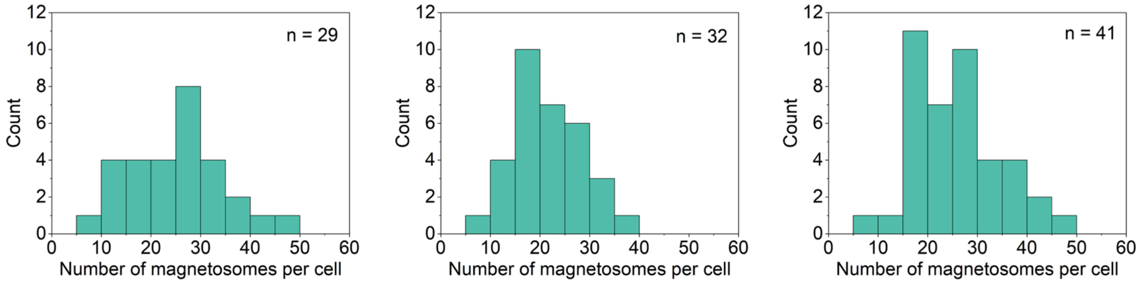


**Supplementary Figure 12. Magnetosome number distribution.** Distribution of the number of magnetosomes per bacterium for the three different bacterial populations used. The amount of magnetosomes was counted only for bacteria totally within the field of view of the micrographs and with clear magnetosome chains.

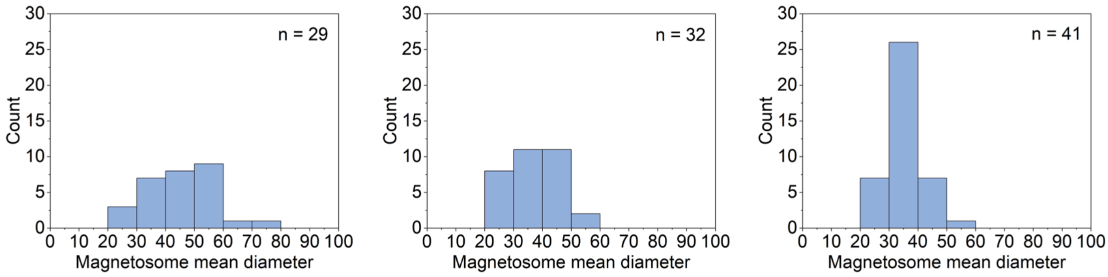


**Supplementary Figure 13. Magnetosome size distribution.** Distribution of the mean size (diameter) of the magnetosomes of one cell for the same bacteria as analyzed in Supplementary Figure 12 for the three different bacterial populations used.

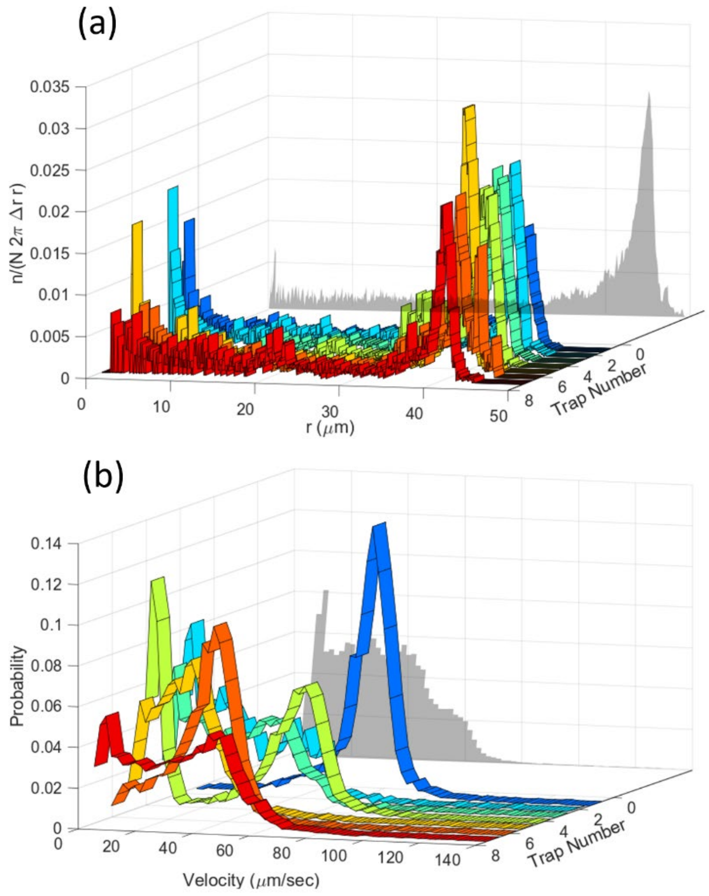


**Supplementary Figure 14: The importance of single-cell analysis.** (a) Experimental radial distributions and (b) corresponding velocity distributions of bacteria in traps of radius $\mathbf{45\ \mu m}$ and for a magnetic field of 500 $\mathbf{\mu}$T. The colored curves show the distributions for individual bacteria in different traps, the gray projected diagram is the averaged behavior of the bacteria population. Averaging over different bacteria can remove important information. This study shows that single-cell analysis is crucial for a quantitative understanding of cell behavior.

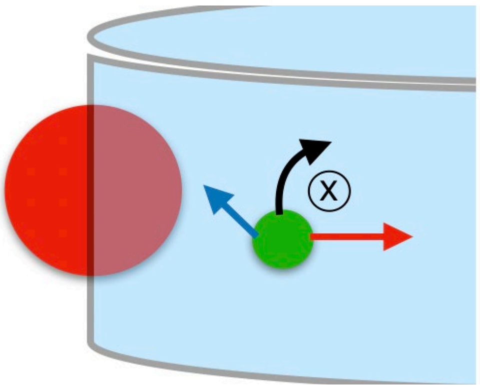

**Supplementary Figure 15. Theoretical wall interaction scheme.** In the simulation, an imaginary sphere of radius $\boldsymbol{A_w}$ (red) is taken to be situated with its center on the wall. The imaginary sphere radius is $\boldsymbol{A_w}$. The bacterium is represented as a green sphere, and its velocity is depicted as a blue arrow; in dark red, the steric repulsion; in black, the hydrodynamic torque.

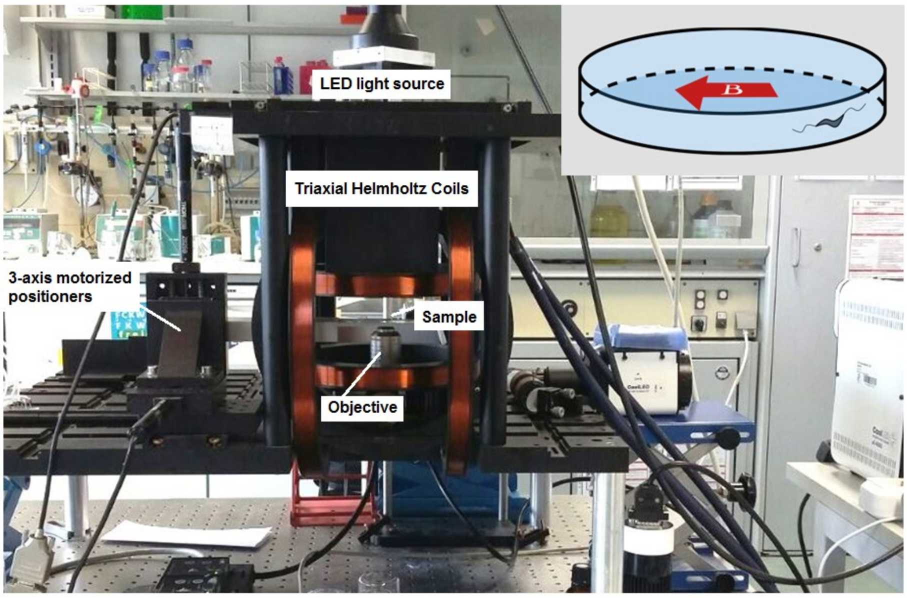


**Supplementary Figure 16: Magnetic microscope**. An inverted optical microscope to take images using a long working distance 40x objective lens. A white light LED was used as a light source to illuminate the sample. The microscope was setup inside 3D-axis Helmholtz coils with a controller with a precision of ±2.5 µT. It was also equipped with three motorized linear stages to provide sample positioning in 3D. **Inset:** The direction of the applied magnetic field in experiments; the magnetic field was set to 0 µT, 50 µT or 500 µT parallel to the microtrap diameter.

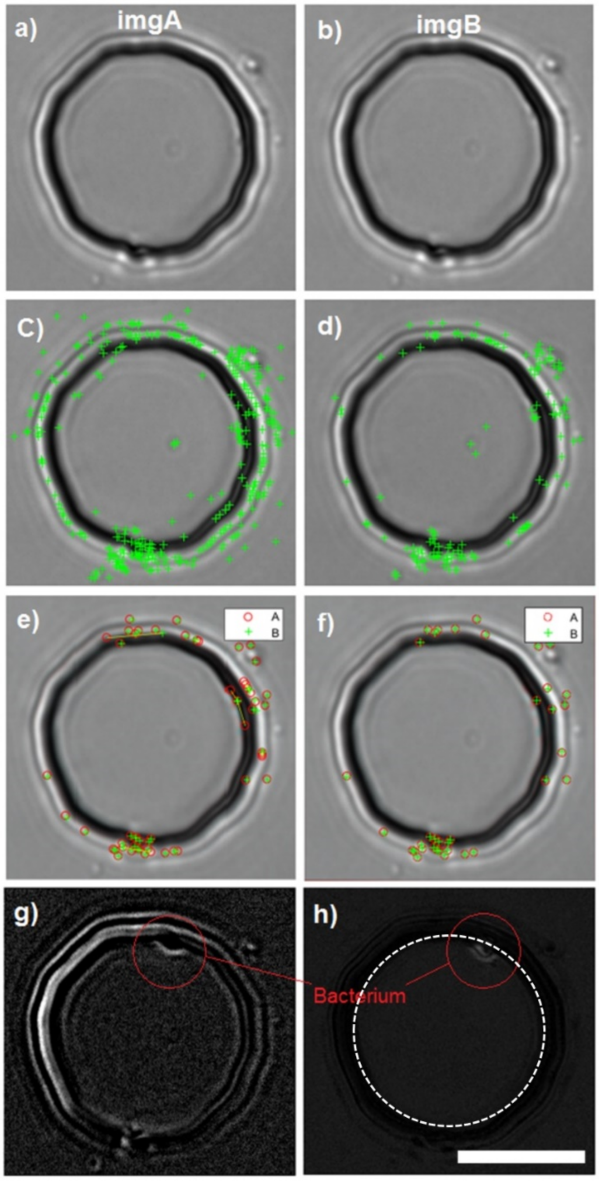


**Supplementary Figure 17: Image stabilization process for effective background subtraction and bacteria tracking**. a) and b) The original successive frames imgA and imgB. c) and d) Detected corners and features that were sharp and changed drastically (the green crosses) in imgA and B, respectively. e) Corresponding points in imgA (red circles) and imagB (green crosses), found according to previously extracted features. f) Apply transformations including scaling, rotation, translation, and shearing on imgB to make the corresponding points points in imgA and imgB coincide the corresponding. g) and h) Result of background subtraction before and after image stabilization, respectively. Dashed white circle in h) is the mean radius of the trap based on the confocal measurements. Scale bar is 10 m.

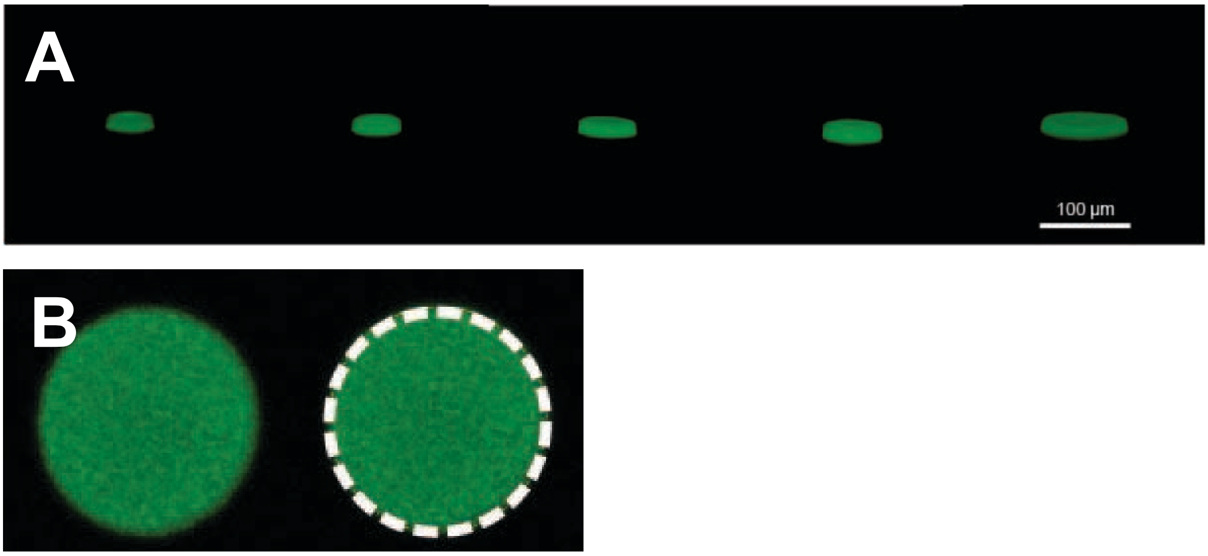


**Supplementary Figure 18. 3D trap reconstruction.** A) 3D reconstructions of the microtraps, from left to right with radii 7.5, 12.5, 17.5, 25, and 45 $\mu m$. The microtraps were filled with calcein and imaged with a confocal microscope. The 3D images were reconstructed with Fiji's 3D viewer plugin (ImageJ). B) x-y plane view used to determine the diameter of microtraps: a circle was fitted to fluorescent area of one microtrap slice. The image corresponds to a microtrap of radius 12.5 $\mu m$.

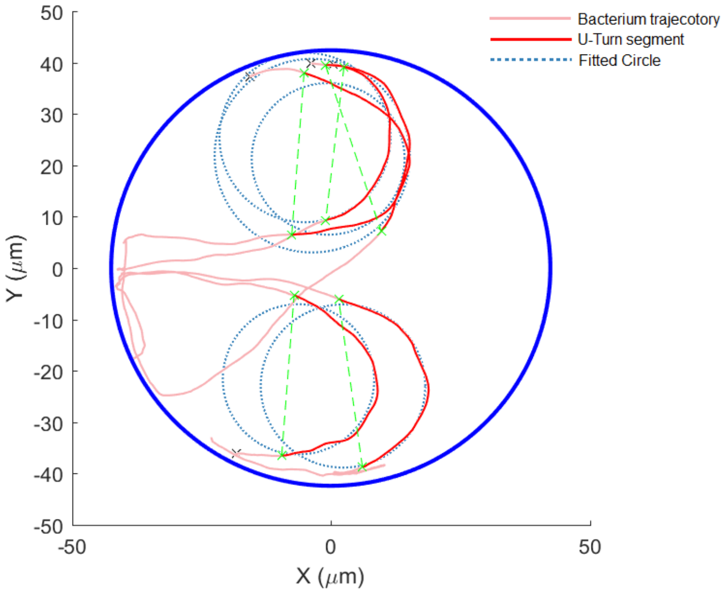


**Supplementary Figure 19. Sample U-turn of bacteria in the micro-trap**. A circle is fitted (dotted blue circles) to the trajectory of the bacterium (light-red trajectories) to extract the radius of the U-turn. The U-turn segment (dark-red segments) is selected for further analysis such as velocity distributions during the U-turn.

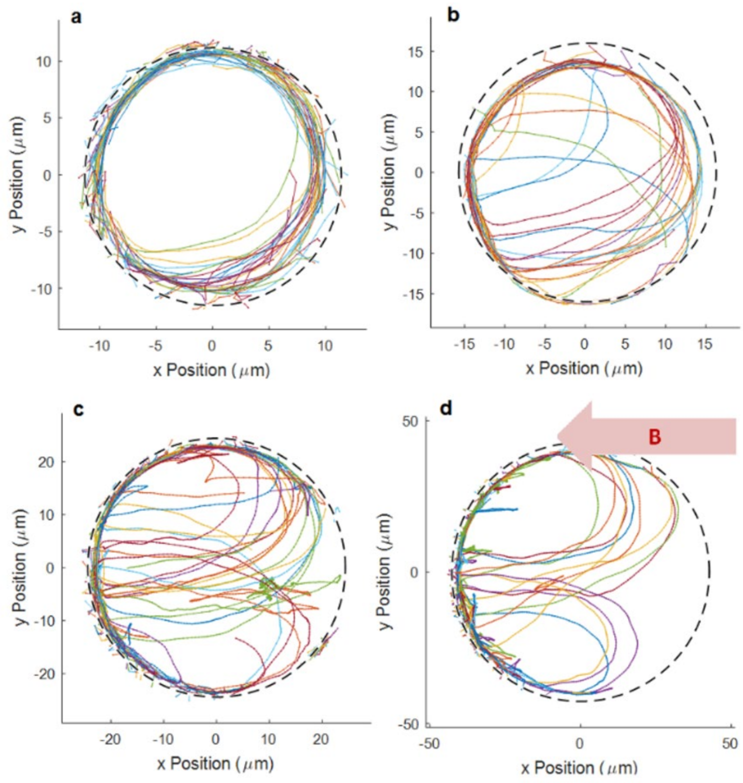


**Supplementary Figure 20: Sample trajectory of bacteria and U-turn.** (a-d) Sample trajectories of bacteria showing U-turns for different microtrap sizes with radii of 12.5, 17.5, 25 and 45 µm, respectively, in the presence of a 500 µT magnetic field. No U-turns were observed in the smallest microtrap with radius of 7.5 µm. Trajectories with different colors correspond to different segments of the bacterium's whole trajectory. The segments arise, because the tracking algorithm sometimes misses points in the trajectories when bacteria swim close to the wall.

**Supplementary Table 1. Measured microtrap diameter.** Extracted mean microtrap diameters from fitting a circle to the fluorescent area of the microtrap x-y slice. The error is calculated by the standard deviation of the measured diameter over different z-stacks and different examined microtraps (n=11 for each microtrap size). The measured values are in good agreement with the nominal values of the printed mask patterns.

| | **Size 1 ($\mu m$)** | **Size 2 ($\mu m$)** | **Size 3 ($\mu m$)** | **Size 4 ($\mu m$)** | **Size 5 ($\mu m$)** |
|---|---|---|---|---|---|
| **Diameter** | **15** | **25** | **35** | **50** | **90** |
| **$D_{measured}$** | 14.85 | 26.52 | 35.8 | 52.13 | 90.88 |
| **$D_{error}$** | 1.04 | 0.44 | 0.42 | 0.65 | 1.05 |

**Supplementary Table 2. Experimental peak positions at 0 magnetic field.** To determine the peak position, the experimental mean distributions at 0 magnetic field (without the long tail) was fitted with the Matlab fitting tool with a gaussian $aexp(-(r-b)^2/c^2)$. The corresponding peak values are reported here.

| Trap size | Trap radius (µm) | Peak position from centre, $b$ (µm) | Sigma, $c/\sqrt{(2)}$ (µm) |
|---|---|---|---|
| **1** | 7.3 | 5.1 | 0.5 |
| **2** | 12.8 | 10.0 | 0.8 |
| **3** | 17.9 | 14.3 | 0.9 |
| **4** | 26.1 | 22.5 | 0.6 |
| **5** | 45.4 | 40.3 | 0.8 |

**Supplementary Table 3. Simulation parameters.** The trap sizes are taken from the experimental measurements with fluorescent microscopy (see Supplementary Table 3). The velocities are obtained from the mean experimental values with no magnetic fields (see Supplementary Figure 8). The hydrodynamic parameters $T_w$ and $A_w$ are obtained by choosing the set of parameters for which the adjusted R squared ($R^2_{adj}$) value between simulated and experimental data is minimum (see the Methods section in the Main Text). The errors (indicated as $\Delta^-$ and $\Delta^+$ respectively for the lower and upper bound) are obtained varying one parameter at the time (keeping the other constant), calculating the $R^2_{adj}$, and allowing it to be 10% less than the maximum. The error on the error is of 0.05 units.

| Size | Trap radius (µm) | Velocity (µm s$^{-1}$) | $T_w$ (adim.) | $\Delta^-_{Tw}$ | $\Delta^+_{Tw}$ | $A_w$ ($\mu m$) | $\Delta^-_{Aw}$ ($\mu m$) | $\Delta^+_{Aw}$ ($\mu m$) | $R^2_{adj}$ |
|---|---|---|---|---|---|---|---|---|---|
| **1** | 7.3 | 30 | 14 | -1 | 1 | 2 | -1 | 1 | 0.70242 |
| **2** | 12.8 | 40 | 11 | -1 | 1 | 3 | -1 | 1 | 0.50831 |
| **3** | 17.9 | 45 | 9.5 | -1.4 | 1.1 | 3.5 | -0.7 | 0.3 | 0.83811 |
| **4** | 26.1 | 30 | 4.5 | -1 | 1.9 | 3.7 | -0.3 | 0.2 | 0.89411 |
| **5** | 45.4 | 40 | 2.8 | -0.7 | 1.1 | 6 | -0.4 | 0.3 | 0.89615 |

**Supplementary Table 4. Master mould fabrication parameters.** SU8 3010 spin-coating, baking and UV exposure parameters used for the fabrication of the master moulds.

| **CONTROL LAYER** (final height: 20 µm) | |
|---|---|
| Spin-coating | 15 s 500 rpm, 30 s 925 rpm |
| Softbake | 3 min 65 °C, 10 min 95 °C, 1 min 65 °C |
| Exposure | 5 s |
| Post-exposure bake | 1 min 65 °C, 6 min 95 °C, 1 min 65 °C |
| **FLUIDIC LAYER** (final height: 10 µm + 10 µm) | |
| Spin-coating | 15 s 500 rpm, 30 s 3,000 rpm |
| Softbake | 1 min 65 °C, 3 min 95 °C, 1 min 65 °C |
| Exposure | 5 s |
| Post-exposure bake | 1 min 65 °C, 3 min 95 °C, 1 min 65 °C |
| Spin-coating | 15 s 500 rpm, 30 s 4,000 rpm |
| Softbake | 1 min 65 °C, 3 min 95 °C, 1 min 65 °C |
| Exposure | 5 s |
| Post-exposure bake | 1 min 65 °C, 3 min 95 °C, 1 min 65 °C |

**Supplementary Video 1.** Real time magnetotactic bacteria (MSR-1) swimming in a micro-trap with radius of 45 m in the absence of external magnetic field. Imaged with 40x objective lens (NA=0.6, air, Nikon) and a sCMOS camera (2560×2160 pixels; Zyla, Andor Technology) with the rate of 50 fps.

**Supplementary Video 2.** Real time magnetotactic bacteria (MSR-1) swimming in a micro-trap with radius of 45 $\mu$m in the presence of external magnetic field (500 $\mu$T, horizontally, north pole directing to left). Imaged with 40x objective lens (NA=0.6, air, Nikon) and a sCMOS camera (2560×2160 pixels; Zyla, Andor Technology) with the rate of 50 fps.
